# Microfabricated disk technology: rapid scale up in midbrain organoid generation

**DOI:** 10.1101/2021.05.31.446446

**Authors:** Nguyen-Vi Mohamed, Paula Lépine, María Lacalle-Aurioles, Julien Sirois, Meghna Mathur, Wolfgang Reintsch, Lenore K. Beitel, Edward A. Fon, Thomas M. Durcan

## Abstract

By providing a three-dimensional *in vitro* culture system with key features of the substantia nigra region in the brain, 3D neuronal organoids derived from human induced pluripotent stem cells (iPSCs) provide living neuronal tissue resembling the midbrain region of the brain. However, a major limitation of conventional brain organoid culture is that it is often labor-intensive, requiring highly specialized personnel for moderate throughput. Additionally, the methods published for long-term cultures require time-consuming maintenance to generate brain organoids in large numbers. With the increasing need for human midbrain organoids (hMOs) to better understand and model Parkinson’s disease (PD) in a dish, there is a need to implement new workflows and methods to both generate and maintain hMOs, while minimizing batch to batch variation. In this study, we developed a method with microfabricated disks to scale up the generation of hMOs. This opens up the possibility to generate larger numbers of hMOs, in a manner that minimizes the amount of labor required, while decreasing variability and maintaining the viability of these hMOs over time. Taken together, producing hMOs in this manner opens up the potential for these to be used to further PD studies.

## 1. Introduction

Parkinson’s disease (PD) is a neurodegenerative disorder, affecting more than 1% of the population over 65 years of age [1]. Classical hallmarks of PD are the loss of dopaminergic (DA) neurons in the substantia nigra pars compacta, accompanied by the presence of neuronal inclusions termed “Lewy bodies”. To date, treatment of PD is limited to symptom management, without any cure to halt the progression of the disease [2]. Thus, it is imperative we better refine the models we use in fundamental research to better understand the pathophysiology of PD and for developing more effective therapeutic strategies.

The brain is the most complex organ in the human body, rendering fundamental research and drug development for neurodegenerative disease extremely challenging. Moreover, the inaccessibility of neuronal tissue from the brain has led to an overreliance on immortalized cancer cells or animal models to capture the architecture and interconnected nature of the brain. Yet, this approach is limited, and often intrinsic differences exist between human neurons and cell types from other species making it difficult to interpret findings. When researchers can access human brain tissue, it is often from autopsies with extensive cell loss, or from surgical samples, meaning that material is limited and often linked to a given disease [3, 4]. Thus, to truly understand the brain, we need to access human neuronal tissue in quantities sufficient for large-scale discovery work.

With the discovery of the Yamanaka factors in 2006 [5], the ability to generate induced pluripotent stem cells (iPSC) from the blood of any individual opened up infinite possibilities to generate customizable and physiological human cell types; and miniaturized organs in a dish. In particular, we can now generate human neurons at levels that had proved elusive. With their self-renewal abilities and potential to be differentiated into disease-relevant cells, iPSCs provide a unique platform to study PD within disease relevant human cell types [6–14], without the difficulties of obtaining neurons from a human surgical sample. Building on these discoveries, novel brain organoid methodologies emerged in 2013 and 2014 from the work of Madeline Lancaster and Juergen A. Knoblich [15, 16]. Since these seminal discoveries, 3D neuronal structures have been used to model early brain development, neurodevelopmental and neurodegenerative disorders, to develop treatments against Zika virus [17–23], and for modelling COVID-19 infection [24–27]. Yet often the methods to generate brain organoids are quite labor intensive, costly and time consuming, limiting their widespread use where large amounts of cells or tissue are required. Often, research groups lack the means to make these organoids at levels needed for translational or fundamental discovery purposes [28].

High throughput screening is also an application which requires a large number of organoids of a consistent quality, and a number of recent publications have pointed the way towards incorporating brain organoids into small molecule screens. In 2021, JC Park and colleagues performed a small molecule screen with 1,300 cerebral organoids generated in AggreWell™ plates, from the iPSCs of healthy participants or sporadic Alzheimer’s disease patients [29]. These plates could then be transferred into 96-well plates for screening purposes. Along these same lines, Renner and others developed an automated workflow to produce thousands of brain organoids in 96-well plates (midbrains and forebrains), from cell seeding to maintenance and analysis using robotic liquid handlers [30]. Such standardized methods reduce the variability of manual handling and demonstrate that methodologies can be applied to grow human brain organoids at large enough numbers when needed.

In our group, with the need for more than 500 human midbrain organoids (hMOs) per batch, we developed and refined our earlier process [7] of producing hMOs to ensure large numbers could be generated from batch to batch with reproducible quality. Our objective was to ameliorate hMO production, making it robust, consistent and scalable, to levels where organoids can be provided for multiple uses across a given batch. In this manuscript we describe our standardized methods to produce hMOs derived from several iPSC lines, with the use of a microfabricated embryoid body disk device (EB disk). We show that large numbers of hMOs can be simultaneously generated, similar in size, morphology, and cellular number. In addition, we describe our workflow for maintaining these hMOs in long-term culture and provide details on our microscopy analysis tools to quantify the area of hMOs and to investigate the 3D structure and total cell numbers of hMOs. With the advent of iPSCs [5] and 3D brain organoids [3, 6–15, 17–21, 31–43], the opportunity is now available to all researchers to work with living neuronal tissue that more closely resembles what we could obtain from the same region in the human brain.

## 2. Material and methods

### 2.1. Generation of hMOs using eNUVIO EB-Disks

#### 2.1.1. List of material, reagents and equipment for generation of hMOs using eNUVIO EB-Disks

A method to generate hMOs using ultra-low attachment 96-well plates was previously described [7]. Background information regarding the different media used to culture hMOs are described in [7]. The media and biochemicals used to generate the results are listed in **Table 1**. Consumables and equipment used are shown in **Table 2**.

**Table 1.**
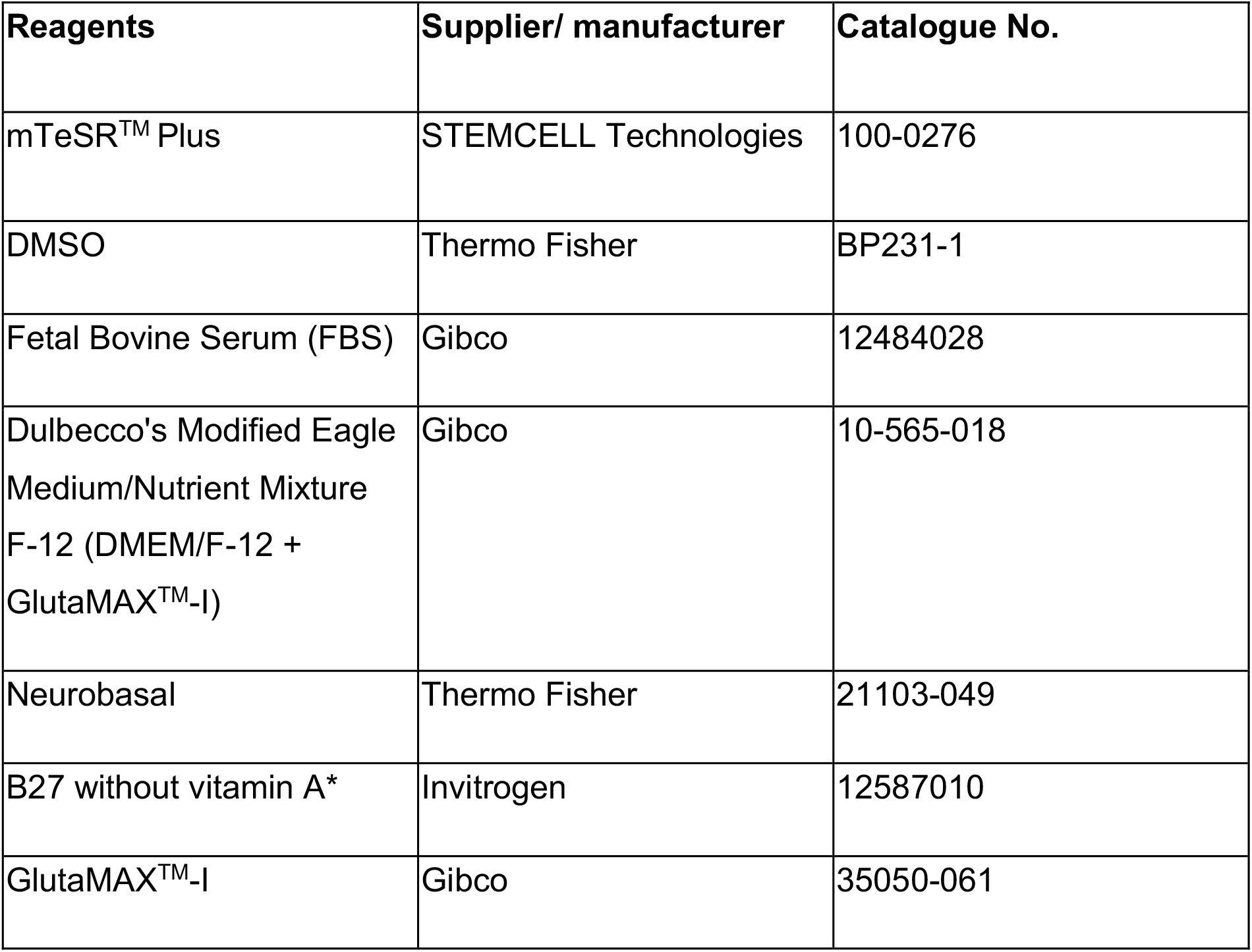
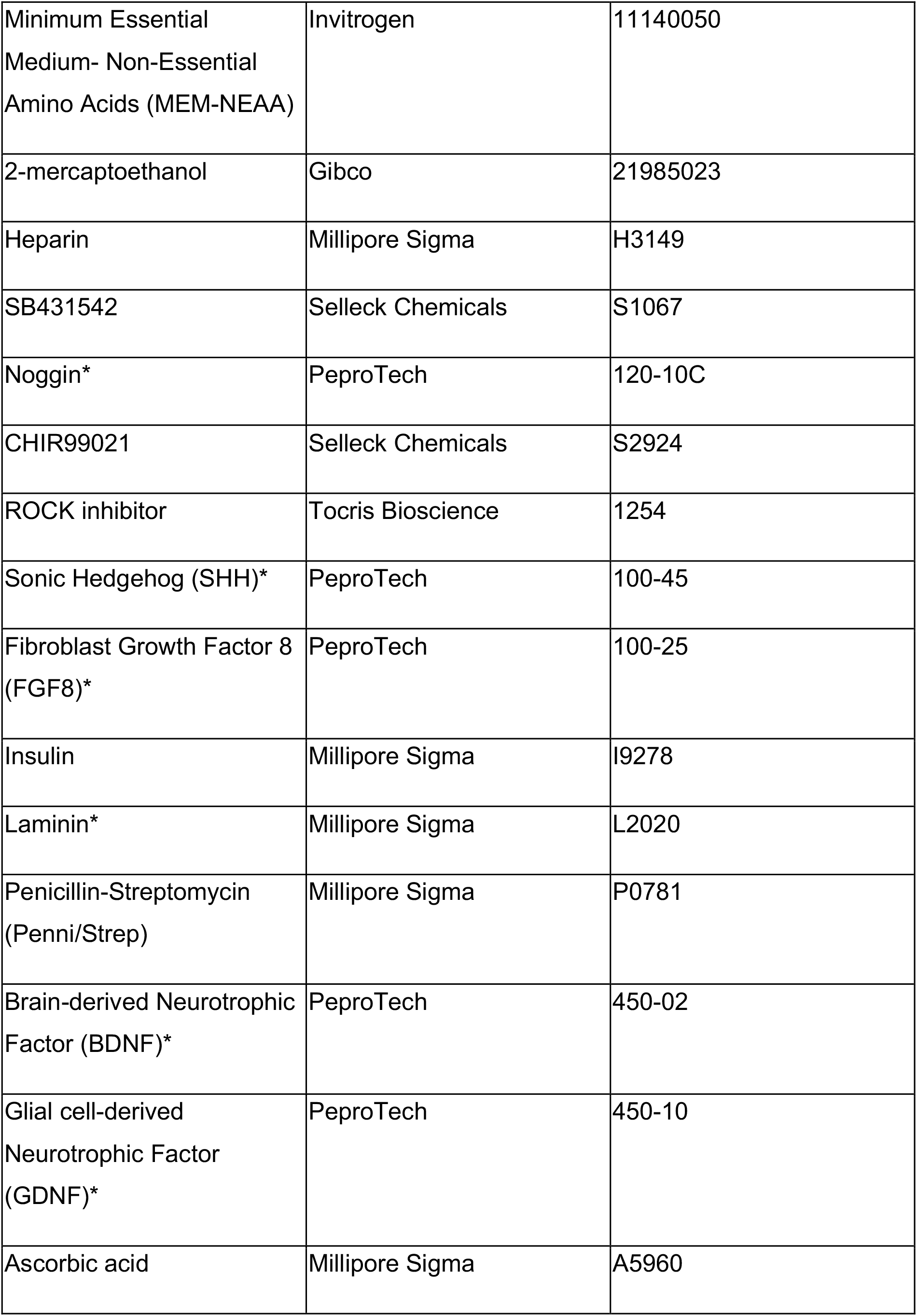
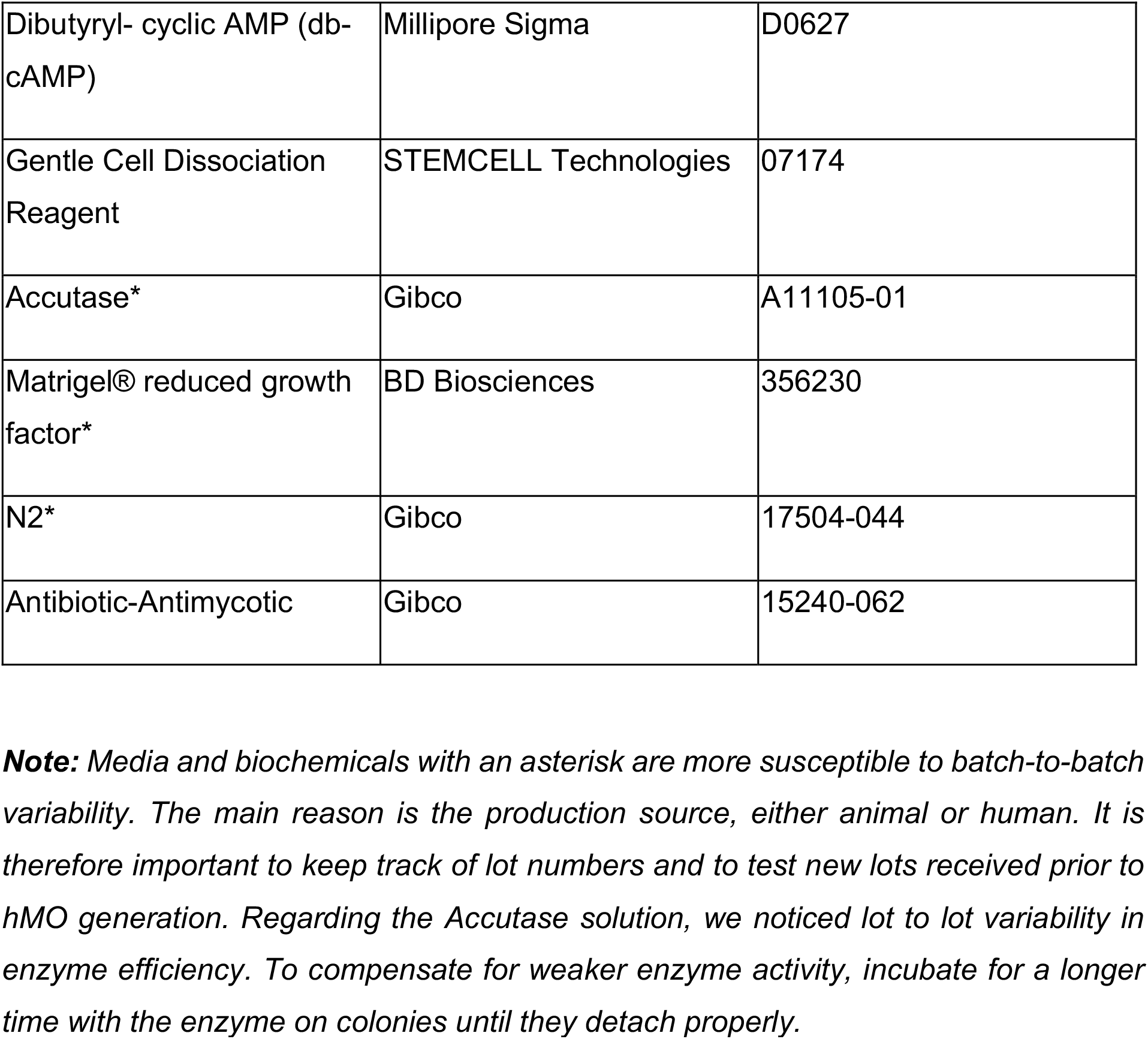
List of media and biochemicals

**Table 2.** List of consumables and equipment

| Consumables and equipment | Supplier | Catalogue No. |
| --- | --- | --- |
| Cell culture dishes, 10 cm | Thermo Fisher | 08772E |
| 5 ml serological pipettes, wrapped | Thermo Fisher | 1367811D |
| EB-Disk<sup>948</sup> ultra low attachment surface | eNUVIO | enuvio.com |
| EB-Disk<sup>948</sup> Centrifuge Adapter (double) | eNUVIO | enuvio.com |
| Cell culture dish, 60 mm | Greiner Bio-One | 628160 |
| Cell culture dish, 150 mm | Falcon | 353025 |
| 15 ml Falcon Conical tube | Thermo Fisher | 352097 |
| 50 ml Falcon Conical tube | Thermo Fisher | 352098 |
| 10 µl ultrafine long tips (autoclave before use in cell culture) | Harvard Apparatus | DV-P1096-FR |
| 100-250 µl non-sterile ultrafine tips, refill package (autoclave before use in cell culture) | VWR | 89368-954 |
| 100-1250 µl non-sterile ultrafine tips, refill package (autoclave before use in cell culture) | VWR | 89079-470 |
| Nitrile gloves | Diamed | TECNITE-SPF |
| Pipette set | Gilson | F167380 |
| Pipette Aid, Drummond | VWR | 53498-105 |
| Centrifuge for 96 well plates | Eppendorf | 022625501 |
| Multichannel pipette | Gilson | FA10015 |
| 50 mL reagent reservoir | Corning | 4870 |
| Box of 1 mL cut tips autoclaved | na | na |
| Metal forceps autoclaved | Thermo Fisher | 12-000-131 |
| Magnetic stirrer, nine position | Chemglass Life Sciences | CLS-4100-09 |
| Reusable glass spinner flask “bioreactor” | Corning® | 4500-500 |
| 37°C incubator | Thermo Fisher | HERAcell 150i |

#### 2.1.2. iPSC lines for generation of hMOs using eNUVIO EB-Disks

To generate hMOs from human iPSCs, it is critical to use cells with minimal differentiation (<5% of iPSC cultures should be non-pluripotent cells), which will reduce organoid variability and increase their quality. The results described were obtained using the iPSC lines listed below. For iPSC culture and maintenance please refer to [7, 44].

##### iPSC lines

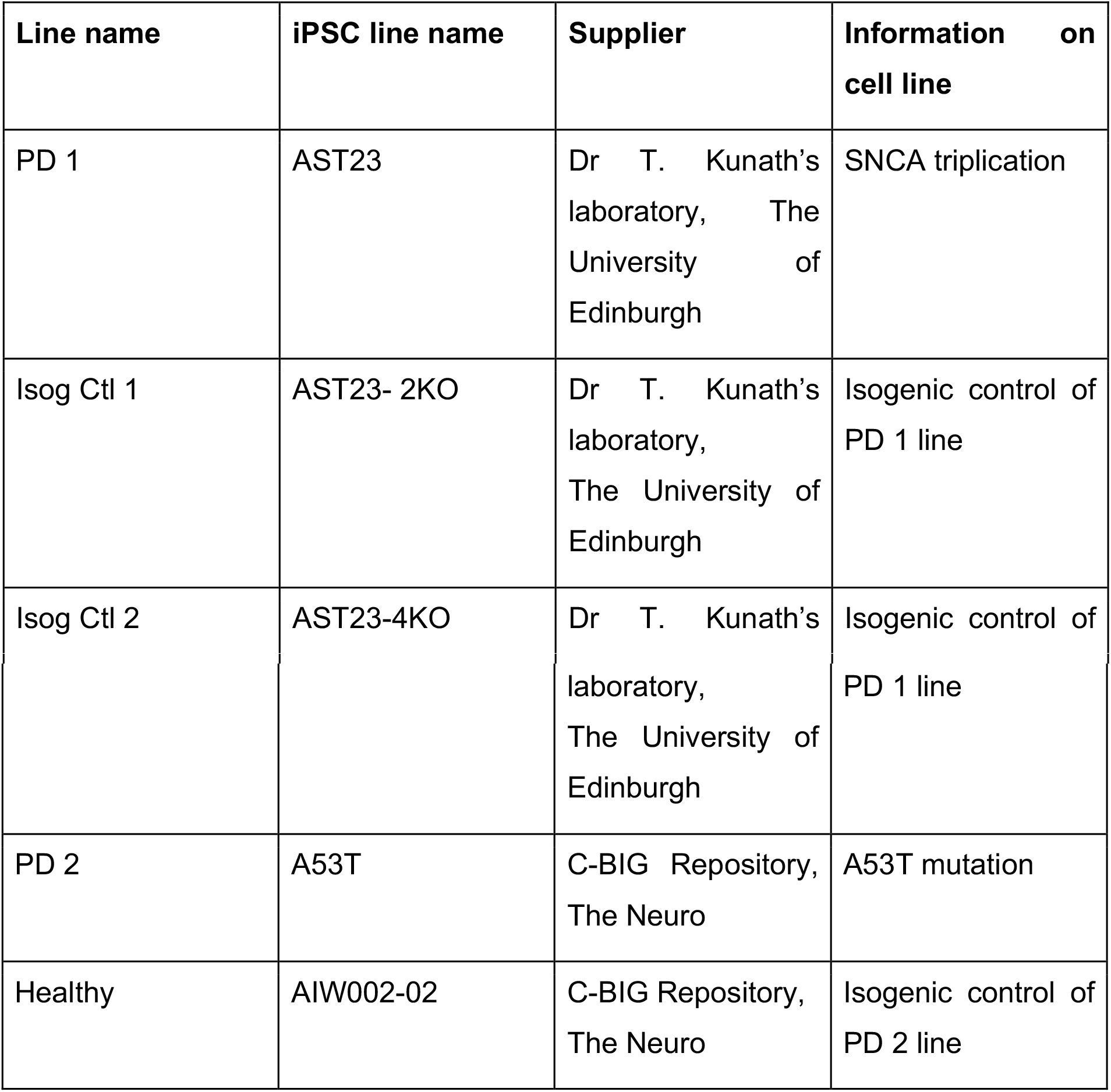

#### 2.1.3. Methods for generation of hMOs from human iPSC using EB-Disk from eNUVIO

In this section we describe the methodology to generate hMOs from a single EB-Disk containing 948 microwells. Starting from iPSC colonies, we describe their dissociation into a single suspension, seeding of the cells into the EB-disk, and the subsequent steps to drive the 3D cellular aggregates toward a midbrain fate until the hMOs are transferred into bioreactors for long-term culture maintenance (**Fig. 1**). Details about the EB-Disk device, timeline, patterning factors and results expected are described in section ***3. Results***.

**Fig. 1.**
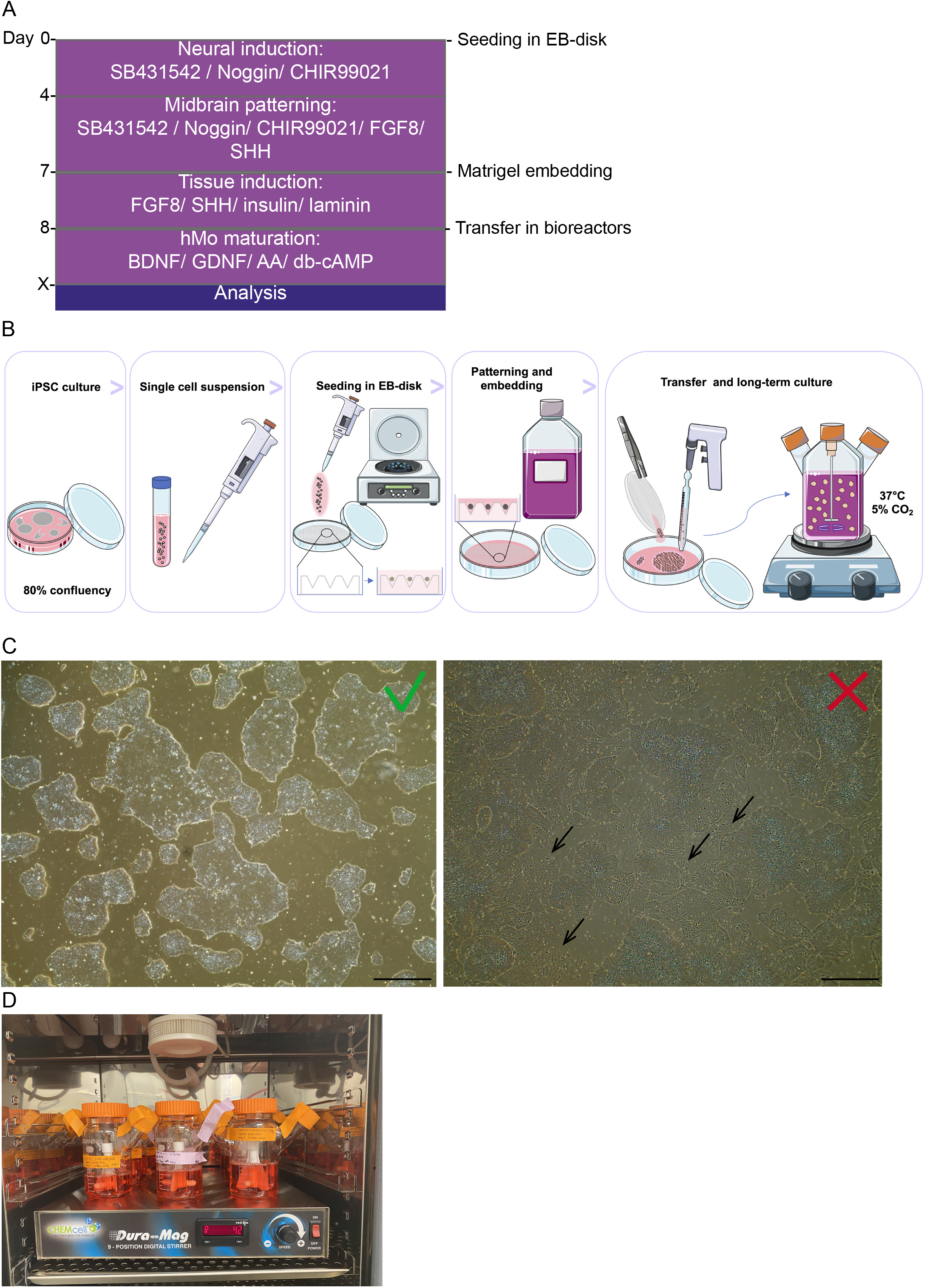
Tools to generate hMOs with an EB-Disk device. (A) Summary of the timeline and patterning factors used to differentiate hMOs from iPSC lines. (B) Schematic for hMO generation from iPSC colonies to long-term culture in a spinning flask. (C) Quality of iPSCs suitable for hMOs formation. The colonies do not present signs of differentiation and reached 80% of confluency. Scale bar= 200 μm. (D) Typical incubator setting with multiple (nine) bioreactors positioned on the magnetic stirrer.

##### Day 0: Seeding of iPSCs

- Start with a 10 cm dish of human iPSCs at 80% confluency. There should be less than 5% differentiated cells. All areas of differentiation must be removed before proceeding to the next steps (**Fig. 1C**).
- Wash cells with 5 mL of DMEM/F-12 + GlutaMAX^TM^-I + Antibiotic-Antimycotic.
- Dissociate cells with Accutase: remove medium used to wash cells and add 5 mL of Accutase to the dish. Incubate at 37°C for 3 min. The colonies should have detached. Then add 5 mL of DMEM/F-12 + GlutaMAX^TM^-I + Antibiotic-Antimycotic to stop the enzymatic reaction.
- Transfer cells to a 15 mL tube, then wash the dish with 5 mL of DMEM/F-12 + GlutaMAX^TM^-I + Antibiotic-Antimycotic to collect the remaining detached cells. Transfer the wash medium to the 15 mL tube.
- Centrifuge for 3 min at 1200 rpm.
- Remove supernatant without disturbing the pellet.
- Add 1 mL of neuronal induction media and resuspend the pellet up and down, using a P1000 pipette with a 1000 μl tip and a 200 μl tip combined.
- Count cells and evaluate the cell viability with trypan blue, which is typically around 95%.
- Prepare a mix of 9.5 million cells in 1.5 mL of neuronal induction media with ROCK inhibitor.
- Add slowly the 1.5 mL cell suspension on top of the EB-Disk^948^ and close the 60 mm dish.
- Place the dish into the centrifuge adapter and centrifuge for 10 min at 1200 rpm.
- Remove the dish from the centrifuge adapter and complete with 5.5 mL of neuronal induction media with ROCK inhibitor. <u>**Note**</u>: the complementary media should be added slowly on the side of the dish to avoid disturbing the aggregates formed in the microwells. The media should fill the dish and properly cover the surface of the EB-Disk.
- Place the 60 mm dish into a 150 mm dish before moving it into the incubator. Three 60 mm dishes can be placed into each 150 mm dish. This step prevents incubator contamination from media spills.

###### Critical

The quality of iPSCs is crucial to successfully generate hMOs. Thus, mycoplasma tests should be performed routinely, we suggest once a month, to confirm the absence of contamination before using the iPSCs to generate hMOs. Moreover, colonies should not present with any sign of differentiation (5% differentiation is the maximum) prior to enzymatic detachment (**Fig. 1C**). Additionally, to guarantee the high viability of the iPSC single cell suspension, the cell detachment step should be monitored carefully and stopped once all colonies are detached. Excessive incubation with Accutase will decrease cell viability and interfere with EBs formation.

##### Day 2: Neuronal induction medium change

###### Observation

EBs should have formed properly with a defined smooth border (**Fig. 2B**).

- To change the media, aspirate out the liquid from the side of the dish. Ideally, insert the end of the tip between the EB-Disk and the border of the dish. Do not aspirate from the top of the microwells to avoid suction of the EBs.
- Delicately pour 7 mL of neuronal induction media without ROCK inhibitor on the side of the dish (**Table 3**). **Note:** pouring the liquid on top of the disk will cause EBs to lift.
- Place back the dish in the incubator.

**Fig. 2.**
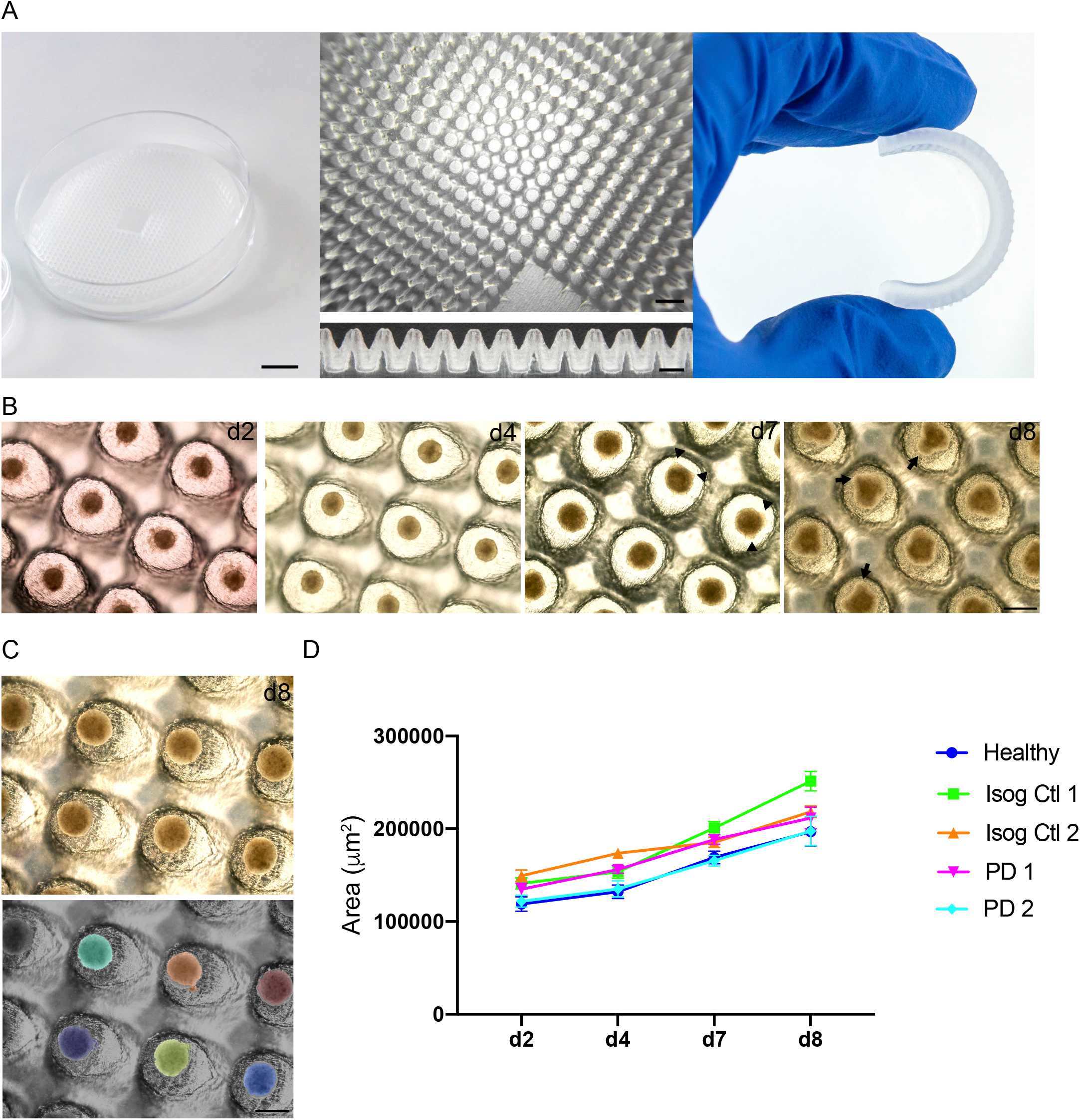
EB-Disk device and EB formation. (A) Microfabricated 948 wells EB-Disk^948^ device. The left picture represents a general view of the EB-Disk^948^ in a dish. Scale bar = 10 mm. The middle top view displays the well array. Scale bar = 3 mm. The side middle view, from a cut disk shows the structure of the microwells. Scale bar=1 mm. The right picture shows the flexibility of the disk. (B) Uniform EBs formed from iPSC colonies were then patterned into neuronal midbrain fate with inductive signals. After 48 hours (d2) in neuronal induction media, EBs presented with smooth edges. On day 4 (d4), the media was changed for midbrain patterning medium. After a further 3 days (d7), EBs started to present neuroectodermal buds (triangles) that became more pronounced after 24 hours in Matrigel embedding (d8, arrows). Scale bar= 600 μm. (C) Quantification of the area of EBs using an in-house CellProfiler macro. As an example, the colored image depicted the surface quantified for each EB from the original upper image (d8, Isog Ctl1 line). Scale bar= 600 μm. (D) Quantification of the EBs areas over time from d2 to d8 in 5 different lines, including PD lines, generated at the same time (n= at least 37 EBs per time point per line)

###### Critical

Considering the ultra-low attachment surface of the disk it is important to not disturb the EBs that form within the wells. Dishes should be handled with care, and not be tilted at any time during the media change. With this in mind, the dish should be kept flat in the biosafety cabinet throughout this step.

##### Day 4: Midbrain patterning medium change

###### Observation

EBs should have grown in size (**Fig. 2B-D**).

- To change the media, aspirate out the liquid from the side of the dish. Ideally insert the end of the tip between the disk and the border of the dish. Do not aspirate on top of the microwells to avoid aggregate suction.
- Delicately pour 7mL of midbrain patterning medium (**Table 3**) on the side of the dish to avoid lifting the EBs up.
- Place back the dish in the incubator.

###### Critical

Considering the ultra-low attachment surface of the disk it is important to not disturb the EBs formed within the wells. Dishes should be handled with care, and not tilted at any time during media change, as described in previous step. Thus, the dish should be kept flat in the biosafety cabinet for this step.

##### Day 7: Embedding and tissue induction medium change

###### Observation

hMOs should have grown and present with some neuroectodermal bud extrusions (**Fig. 2B-triangles**). At this step hMOs attain approximately 500 μm in diameter (**Fig. 2D**).

###### Note

Matrigel® reduced growth factor should be thawed slowly at 4°C. We recommend putting a bottle of Matrigel® at 4°C on Day 6 (d6). Place 5 mL pipettes or 1 mL tips at 4°C at the same time.

- Remove the media by aspirating out the liquid from the side of the dish. Ideally insert the end of the tip between the EB-Disk and the border of the dish. Do not aspirate from the top of the microwells to avoid suction of the aggregates.
- Add 1.5 mL of Matrigel® reduced growth factor with cold tips or a cold pipet on top of the EB-disk.
- Place the dish back into the incubator for 30min.
- After 30 min, under the biosafety cabinet, add 7 mL of tissue induction medium (**Table 3**) from the side of the dish. Place the dish back into the incubator.

###### Critical

1) Considering the ultra-low attachment surface of the disk, it is important to not disturb the hMOs formed within the wells. Dishes should be handled with care, and not be tilted at any time during media change. As a result, the dish should be kept flat in the biosafety cabinet for this step. 2) Matrigel® reduced growth factor is temperature sensitive and gels when the environment is warmer than 4°C, thus it is critical to always keep it on ice when removing it from the fridge, and to manipulate it only with cold tips or cold pipets to prevent it from gelling.

##### Day 8: Transfer into spinner flask bioreactors and final differentiation medium change

###### Observation

hMOs will have grown in size and present with more neuroectodermal buds extruding (**Fig. 2B-arrows**).

###### Note

Metal forceps must be autoclaved before performing this step. In addition, bioreactors must be assembled and autoclaved according to manufacturer’s instructions.

- Using autoclaved metal forceps, delicately transfer the EB-Disk to a 150 mm dish containing 10 mL of final differentiation medium (**Table 3**). Place the disk with the microwells facing downwards.
- Gently agitate the disk with the metal forceps, a number of embedded hMOs should float out into the medium.
- Transfer hMOs with a 10 mL pipet broken at the tip (**Fig. S1A**), to an autoclaved 500 mL spinning bioreactor containing 300 mL of final differentiation media.
- Repeat the two previous steps until no more hMOs remain in the disk or the media.
- Place the bioreactor on a magnetic stirrer set at 42 rpm inside a 37°C incubator with 5% CO_2_. At this stage, hMOs can be observed floating in the media.
- Loosen the lateral caps to ensure proper air-gas exchange.

###### Critical

Matrigel® reduced growth factor embedding provides a scaffold for hMOs to develop at early stages. Use care when transferring these hMOs into bioreactors to preserve the scaffold and not to damage it. The use of 10 mL pipettes, broken at the extremity under the biosafety cabinet, in sterile conditions, is required (**Fig. S1A**). Alternatively, you could use 25 mL and 50 mL pipettes as they have wider tips.

###### Suggestion

In this protocol we did not describe the reuse of the EB disks, but it is possible to rejuvenate the disks by following the manufacturer’s guidelines: https://enuvio.com/tele/2021/02/EB-Disk-User-Guide-v2.1.pdf.

##### Day 15 and following weeks: Weekly maintenance of the spinner flask bioreactors

###### Observation

hMOs have grown to full size within the bioreactors (**Fig. 3A**).

- Close the lateral caps before removing the bioreactors from the incubator.
- To change the media, aspirate out the media from one lateral cap. Be careful to not aspirate the hMOs. Leave 100 mL of media on top of hMOs before pouring in 400 mL of fresh final differentiation media.
- Place the bioreactors back on the magnetic stirrer set at 42 rpm inside the incubator. You should see the hMOs floating in the media with a fluid motion.
- Loosen the lateral caps to ensure a proper air-gas exchange.

**Fig. 3.**
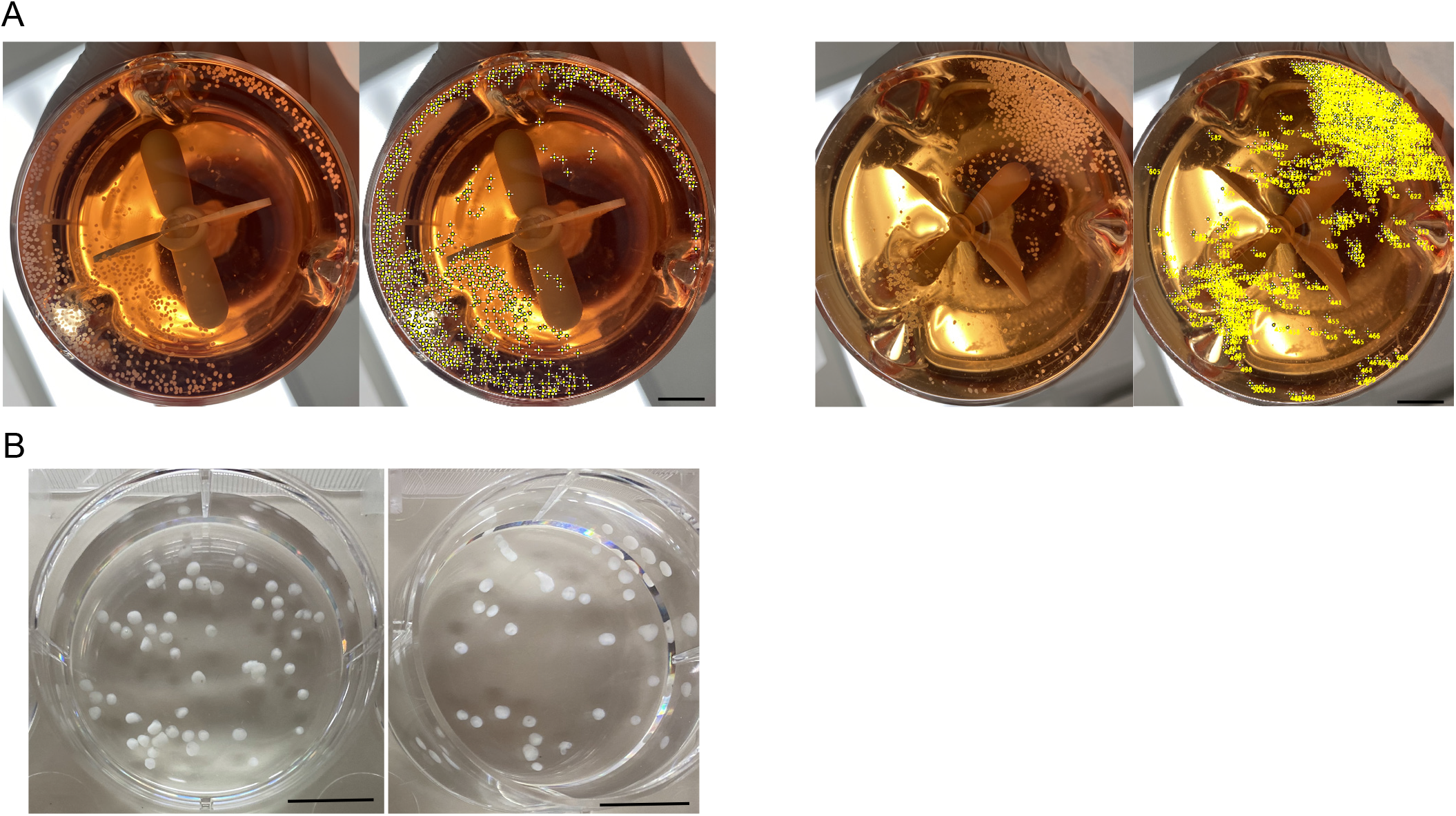
Maturation of hMOs. (A) Quantitation of hMOs after 10 days in final differentiation media. In the left panel, 650 hMOs were counted for hMOs derived from PD 1. In the right panel, 714 hMOs were counted for hMOs derived from Isog Ctl 2. Scale bar= 1 cm. (B) Macroscopic view of 100 days old hMOs derived from Isog Ctl 1 and Healthy line.

###### Critical

Volume of media should be increased up to 500 mL in bioreactors when hMOs become bigger and turn the media yellow after one week. After media change, make sure the bioreactors are properly replaced on the cross marks, on the magnetic stirrer, to ensure the proper motion of the media spin within the flask. A media spin disturbance will lead to the disintegration of the hMOs. Perform mycoplasma test every month to ensure the quality of the material prior analytic experiments.

**Table 3.** Media composition for hMO generation

| **Reagent** | **Neural**<br>**induction**<br>**(50 mL)** | **Midbrain**<br>**patterning**<br>**(50 mL)** | **Tissue**<br>**induction**<br>**(50 mL)** | **Final**<br>**differentiation**<br>**(50 mL)** |
| --- | --- | --- | --- | --- |
| DMEM/-F-12 +<br>GlutaMAX™-I +<br>Antibiotic-Antimycotic<br>/Neurobasal<br>(1:1) | 25 mL +<br>25 mL | 25 mL +<br>25 mL | x | x |
| Neurobasal | x | x | 50 mL | 50 mL |
| 1:100 N2 | 0.5 mL | 0.5 mL | 0.5 mL | 0.5 mL |
| 1:50 B27<br>without vitamin<br>A | 1 mL | 1 mL | 1 mL | 1 mL |
| 1%<br>GlutaMAX™-I | 0.5 mL | 0.5 mL | 0.5 mL | 0.5 mL |
| 1% MEM-NEAA | 0.5 mL | 0.5 mL | 0.5 mL | 0.5 mL |
| 2-mercaptoethano<br>l | 0.175 µL | 0.175 µL | 0.175 µL | 0.175 µL |
| 1 µg/mL Heparin | 50 µL | 50 µL | x | x |
| 10 µM<br>SB431542 | 50 µL | 50 µL | x | x |
| 200 ng/mL<br>Noggin | 50 µL | 50 µL | x | x |
| 0.8 μM<br>CHIR99021 | 13.5 μL | 13.5 μL | x | x |
| 10 μM ROCK<br>inhibitor | 50 μL | x | x | x |
| 100 ng/mL<br>(or 200 ng/mL)<br>SHH | x | 25 μL<br>(or 50 μL) | 25 μL<br>(or 50 μL) | x |
| 100 ng/ mL<br>FGF8 | x | 50 μL | 50 μL | x |
| 2.5 μg/mL<br>insulin | x | x | 12.5 μL | x |
| 200 ng/mL<br>laminin | x | x | 8.5 μL | x |
| 10 ng/mL BDNF | x | x | x | 25 μL |
| 10 ng/mL GDNF | x | x | x | 25 μL |
| 100 μM ascorbic<br>acid | x | x | x | 25 μL |
| 125 μM db-cAMP | x | x | x | 12.5 μL |
| Penni/Strep | x | x | 0.05 mL | 0.05 mL |

### 2.2. In-house CellProfiler pipeline for EB and hMOs area quantification within the EB-Disk

#### 2.2.1. Methods for areas quantification

To measure the size of EBs and hMOs within the disk, brightfield images of dishes containing 4-6 organoids per field of view were analyzed with CellProfiler software (version 4.1.3), www.cellprofiler.org [45]. The analysis pipeline developed is available at this link: https://doi.org/10.17605/OSF.IO/XJAQ3. The following provides a short summary of the analysis steps. First, the brightfield color images, taken with an EVOS XL Core, were transformed to greyscale, inverted to obtain bright organoids with a dark background and smoothened to remove small artefacts. Next, the primary objects (organoids) were identified (using Otsu thresholding). Objects touching the image border were discarded. The size and shape of the primary objects was measured, and a shape filter applied to remove occasional misidentifications in the image. The final organoid masks were overlayed with the original input image for visual control of the organoid identification. Size and shape measurements were exported as CSV files and the average organoid area per image set calculated. The values of area in pixels were then converted to μm^2^ according to the microscope scale.

### 2.3. Flow cytometry analysis of hMOs

#### 2.3.1 List of material, reagents, equipment and software for flow cytometry

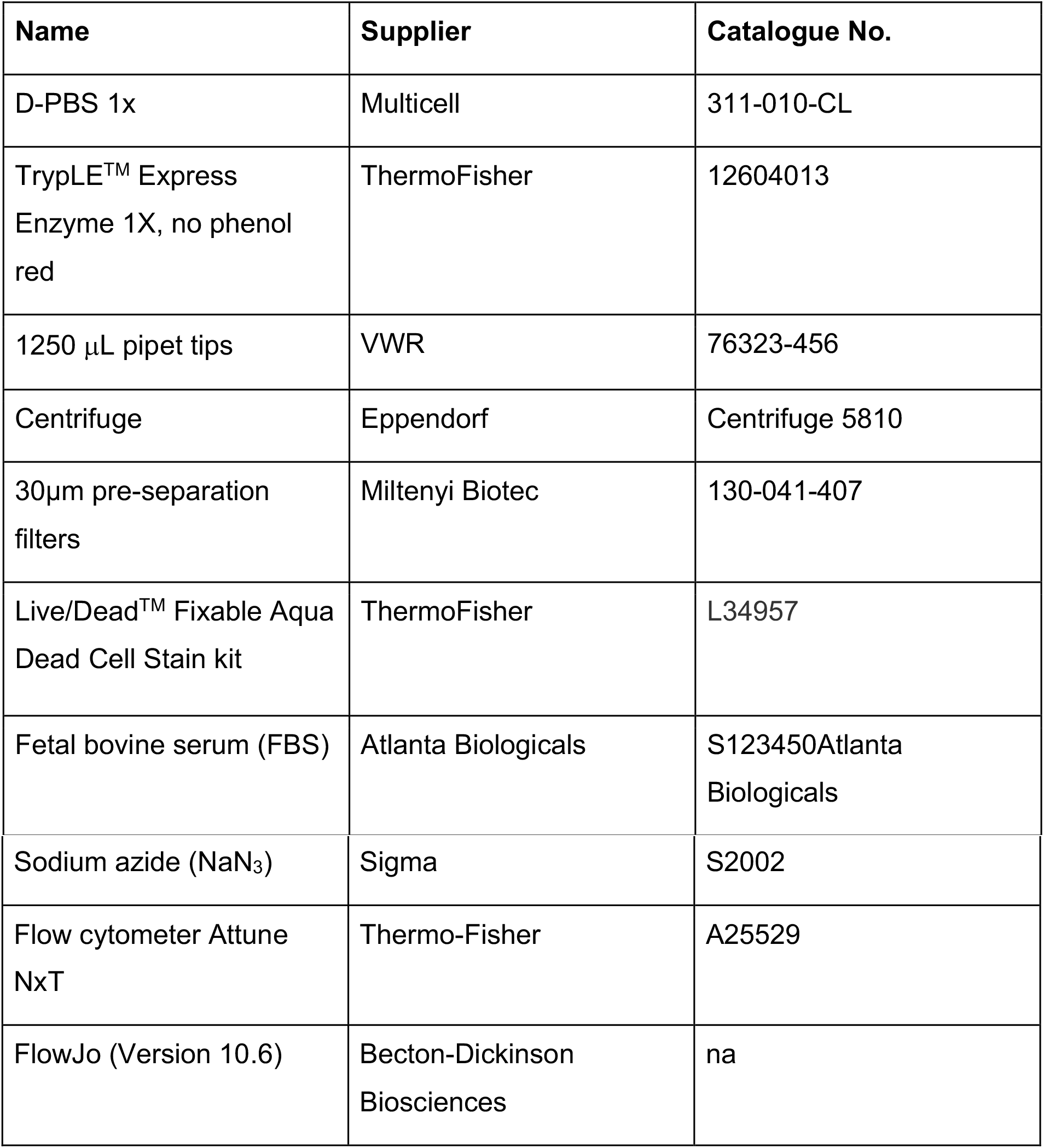

| Name | Supplier | Catalogue No. |
| --- | --- | --- |
| D-PBS 1x | Multicell | 311-010-CL |
| TrypLE™ Express Enzyme 1X, no phenol red | ThermoFisher | 12604013 |
| 1250 $\mu\text{L}$ pipet tips | VWR | 76323-456 |
| Centrifuge | Eppendorf | Centrifuge 5810 |
| 30$\mu\text{m}$ pre-separation filters | Miltenyi Biotec | 130-041-407 |
| Live/Dead™ Fixable Aqua Dead Cell Stain kit | ThermoFisher | L34957 |
| Fetal bovine serum (FBS) | Atlanta Biologicals | S123450Atlanta Biologicals |
| Sodium azide (NaN<sub>3</sub>) | Sigma | S2002 |
| Flow cytometer Attune NxT | Thermo-Fisher | A25529 |
| FlowJo (Version 10.6) | Becton-Dickinson Biosciences | na |

#### 2.3.2. Methods for flow cytometry

First, hMOs were harvested, washed with 1x D-PBS and treated with TrypLE Express before being dissociated with the gentleMAC Octo Dissociator (Miltenyi Biotec) at 37°C to create a single cell suspension. The TrypLE reaction was stopped by adding 1x D-PBS. The single cell suspension was next filtered through a 30 μm mesh (Miltenyi Biotec) and cells were pelleted through centrifugation at 350g for 5 min. Pelleted cells were resuspended in 1x D-PBS to faciliate cell counting on the Attune NxT FACs analyzer (Thermo-Fisher). Following cell counting, at least 200,000 cells were stained with Live/Dead Fixable Aqua (Invitrogen) for 30 min at room temperature (RT) (protected from light). Cells were subsequently washed with 1x D-PBS and centrifuged at 350g for 5 min. The cells were washed twice with FACS Buffer (5% FBS, 0.1% NaN3 in D-PBS) and centrifuged at 350g for 5 min. Cells were then ready for flow cytometry acquisition. For flow cytometry settings and parameters, voltages were set up according to optimal PMT sensitivity using the peak 2 (Spherotech) voltration technique described previously by Maeker and Trotter [46]. All data was acquired on the Attune NxT. Finally, all data generated was analyzed using FlowJo (Version 10.6) (Becton-Dickinson Biosciences).

### 2.4. Fixation and immunofluorescence staining of hMO cryosections

For detailed protocols, please refer to our previous publications [7, 47]. Briefly, hMOs were fixed and cryosectioned according to our published methods [7]. For staining, 14 μm cryosections were rehydrated in phosphate-buffered saline (PBS) for 15 min and surrounded with a hydrophobic barrier drawn with a barrier pen. The sections were then incubated with a blocking solution (5% normal donkey serum, 0.05% BSA, 0.2% Triton X-100 in PBS) for 1 hour at RT in a humidified chamber. The sections were subsequently incubated overnight at 4°C with primary antibodies (S100β 1/200, Sigma S2532; MAP2 1/500, EnCor Biotech CPCA-MAP2; TH 1/200, Pel-Freeze P40101-150) diluted in blocking solution. The following day, cryosections were washed three times in PBS (15 min each) and then incubated in secondary antibodies (donkey antichicken Alexa 488, Jackson ImmunoResearch 703-545-155; donkey anti-mouse Dylight 550, Abcam ab96876, donkey anti-rabbit Dylight 650, Abcam ab96894) diluted in a blocking solution for one hour at RT. Sections were next washed three times in PBS for 15 min each, incubated with Hoechst (1/5000 in PBS) for 10 min, and finally washed once in PBS for 10 min. Finally, sections were mounted (Aqua-Poly/Mount, Polysciences), and the images acquired with a Leica TCS SP8 confocal microscope.

### 2.5. RNA isolation, cDNA synthesis, and qPCR

RNA was purified from iPSCs and hMOs using a NucleoSpin RNA kit (Takara) according to the manufacturer’s instructions. cDNA was generated using iScript Reverse Transcription Supermix (BioRad). Quantitative real-time PCR was performed on a QuantStudio 3 Real-Time PCR System (Applied Biosystems). This produced a total reaction volume of 10 μl, including 5 μl of fast advanced master mix (Thermofisher Scientific), 1 μl 20X Taqman assay (IDT), 1 μl of cDNA and 3 μl of RNAse-free water. Relative gene expression levels were analyzed using the Comparative CT Method (ΔΔCT method). The results were normalized to the GAPDH expression. The relative quantification (RQ) was estimated according to the ΔΔCt methods [48].

### 2.6. Tissue clearing and imaging on hMOs

Despite the small size of hMOs, the three-dimensional (3D) imaging of organoids is challenging due to the intrinsic opacity of the biological tissues. Tissue clearing techniques make biological samples transparent by reducing light scattering and absorption throughout by removing lipids from cell membranes and reducing water content [49]. Here we use an optimized version of the water-based clearing protocol, Clear, Unobstructed Brain Imaging Cocktails (CUBIC) [50, 51] described by Gómez-Gaviro et al. [52]. CUBIC is a non-toxic and straightforward to implement protocol that facilitates antibody penetration during immunolabeling, reduces fluorescence quenching and ensures high tissue transparency for optimal 3D imaging of hMOs.

#### 2.6.1. List of material, reagents and equipment for tissue clearing and imaging

The composition and methods to prepare CUBIC reagent 1 and reagent 2 are described in Gómez-Gaviro et al. [52] and **Table 4**.

**Table 4.**

| **Name** | **Supplier** | **Catalogue No.** |
| --- | --- | --- |
| PBS 1X | Multicell | 311-010-CL |
| Paraformaldehyde | Sigma | 158127-500G |
| Distilled water |  |  |
| Urea | Sigma | 15604 |
| N, N, N', N'-tetrakis (2-hydroxy-propyl) ethylenediamine | Sigma | 122262-1L |
| Triton X-100 | BioShop Canada | TRX777.500 |
| Triethanolamine | Fisher Chemical | T407-500 |
| D-Sucrose | Fisher | BP2020-1 |
| Bovine serum albumin | Multicell | 800-095-CG |
| Sodium azide (NaN<sub>3</sub>) | Sigma | S2002-500G |
| Donkey serum | Sigma | DD9663-10ML |
| Superfrost plus microscope slides | Fisherbrand | 12-550-15 |
| Microscope coverglass | Fisherbrand | 12543D |
| Imaging spacer | SecureSeal™, Electron Microscopy Sciences | 70327-8S |
| Microscope Leica SP8 | Leica Microsystems Inc |  |

#### 2.6.2. Methods for tissue clearing and 3D imaging of hMOs

Live hMOs were rinsed in 1X PBS and fixed for 2 hours at room temperature (RT), washed thoroughly in 1X PBS and incubated in a shaker (80 rpm, 37°C) in CUBIC reagent 1 (R1, for lipid removal) until translucent (~2 days). hMOs were then washed 3 times in 1X PBS at RT (~1 h each) and incubated with blocking solution for 1 hour previous to immunofluorescence (IF) imaging. For IF, hMOs were incubated in a shaker (100 rpm, 37°C) for 3 days with primary antibodies: rabbit anti-tyrosine hydroxylase (TH, 1:500, Pel-Freez Biologicals, AR, USA) and chicken anti-microtubule associated protein 2 (MAP2, 1:500, EnCor Biotechnology Inc, FL, USA), followed by an incubation (3 days) with secondary antibodies: donkey anti-chicken Alexa 488 (1:500, Jackson ImmunoResearch Laboratories, Inc, PA, USA) and donkey anti-rabbit Dylight 650 (1:500, Abcam, MA, USA). hMOs were then washed overnight with 1X PBS at 4°C and incubated with Hoechst 33342 solution in 1X PBS for 2 h (1:1000, Thermo Fisher Scientific) and washed in PBS for 2 h. Immunolabeled hMOs were incubated in CUBIC R2 (for optical clearing) until samples were completely transparent (~1 day).

On the day of imaging, hMOs were placed on a microscope slide carrying four imaging spacers (0.12 mm depth each) piled (SecureSeal™ Imaging Spacers, Electron Microscopy Sciences, PA, USA), filled with CUBIC R2 and covered with a coverglass. hMOs were imaged in a Leica TCS SP8 confocal (Leica Microsystems Inc., ON, Ca) using a HC PL APO 10x/0.40 CS objective, free working distance: 2.2 mm. Image size 512 x 512, voxel size 2.268 x 2.268 x 5 μm (**see Movie**).

#### 2.6.3. 3D image processing and analysis

3D images were processed, and volumes quantified using the free software tool Fiji [53]. For each channel, the background of the images was subtracted with a rolling ball radius =50.0 pixels. The images were binarized with a “default” method, and the volumes quantified with the voxel counter plugin. Data was analyzed and graphically presented using GraphPad Prism (version 8.2.1).

##### Note

hMOs accumulate spontaneous dark neuromelanin granules (**Fig. 4E**) that will create a local black shadow during imaging.

**Fig. 4.**
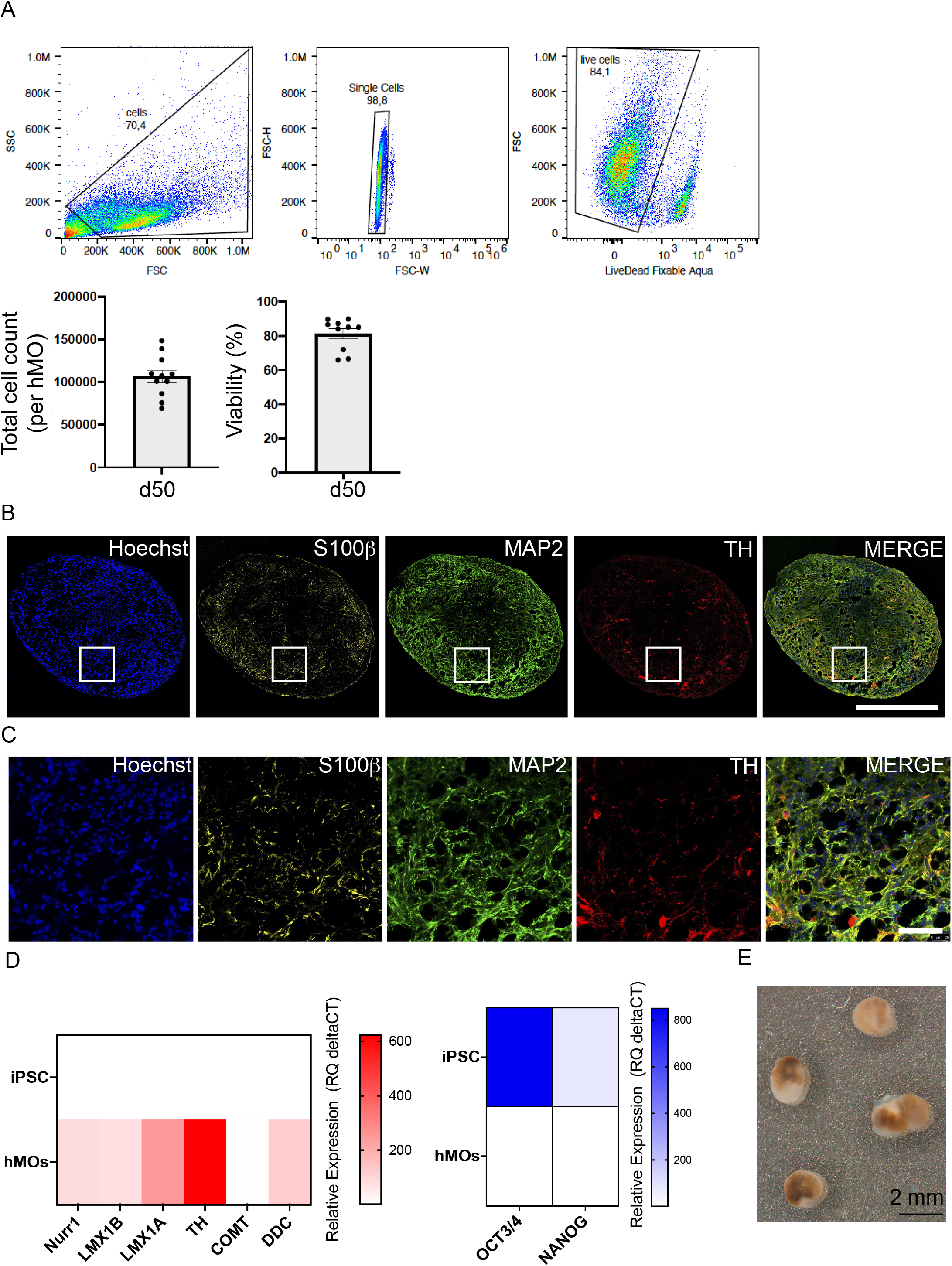
Composition of hMOs. (A) Flow cytometry of 50-day old hMOs to count cells within hMOs after single cell suspension (n=11 Healthy hMOs, mean +/− SEM) and to assess cell viability with Live/Dead staining (n=10 Healthy hMOs, mean +/− SEM). (B) Cryosections of 50-day old Isog Ctl hMOs stained for neurons (MAP2), dopaminergic neurons (TH), astrocytes (S100β) and nuclei (Hoechst). Scale bar = 800 μm. (C) Higher magnification of *(B)* white squares showing neurons (MAP2), dopaminergic neurons (TH), astrocytes (S100β) and nuclei (Hoechst). Scale bar = 50 μm. (D) Realtime PCR depicting normalized expression levels of midbrain RNAs (in red) for Nurr1, LMX1A, LMX1B, TH, COMT, and DDC at 50 days in Isog Ctl hMOs, normalized to endogenous GAPDH and actin controls (n=3, mean). Similar analysis for iPSC transcripts OCT3/4 and NANOG (in blue). (E) Representative image of spontaneous accumulation of neuromelanin granules in Isog Ctl hMOs 170-days old hMOs. Scale bar = 2 mm.

## 3. Results

We derived hMOs from different iPSC lines using classical midbrain patterning factors [14, 54–60], following our previously published timeline [7] (**Fig. 1A**). To generate hMOs from a single EB-Disk containing 948 microwells, we started from iPSC colonies at 80% confluency with no differentiated areas. The process to generate hMOs from iPSCs took eight days. After dissociation of the iPSCs into a single cell suspension, we seeded the cells into one EB-Disk and centrifuged the device to form one cellular aggregate within each microwell of the disk. The subsequent steps consisted of driving the cells toward a midbrain fate with several media changes in stationary culture, until the hMOs were embedded directly on the EB-Disk at day 7 (d7) in a Matrigel® scaffold, to promote the tissue development. Finally on day 8 (d8), the disk was lifted up and flipped to a dish containing final differentiation media. The Matrigel®-embedded hMOs were then transferred into bioreactors containing final differentiation media to promote the maturation of the hMOs and maintained in culture long-term prior their collection at defined time points for experiments (**Fig 1B**). We observed that the condition of the iPSCs is a primary determinant for successfully generating high-quality hMOs. iPSC colonies were maintained daily and passaged in order to present with minimal areas of differentiated cells/cells that had lost their pluripotency (**Fig. 1C**). We used several disks in parallel across different lines, synchronized in confluency, to generate batches with different conditions. The aggregates in each disk were then transferred into a specific bioreactor and placed on a magnetic stirrer housed within an incubator. Each magnetic stirrer can hold up to nine bioreactors (**Fig. 1D**).

The generation process described in this manuscript, to scale-up the production of hMOs, was based on the use of a microfabricated device or “EB-Disk” detailed in **Fig. 2A**. For the production of batches with several hundred hMOs, we used EB-Disks^948^ containing 948 microwells (**Fig. 2A, left and top middle pictures**) pre-coated with an ultra-low attachment coating, composed of 5% Pluronic acid F-127, and with defined dimensions as depicted on the lateral sectioned disk. The diameter of each disk was 52 mm and could be fitted into a 60 mm round culture dish. The volume of each microwell was 1 mm^3^ while the microwell diameter at the base was 900 μm. The disks were made from PDMS, a flexible material, permitting easy handling of the platform for hMO production, while being biologically compatible with the generation of neuronal tissue [61, 62].

We observed the progressive appearance of aggregates within the disk from the seeding day (d0) until their transfer into bioreactors (d8). Indeed, after 48 hours in the disk, EBs formed and presented with smooth edges (**Fig. 2B**). They grew and started to present with neuroectodermal buds after 3 days in midbrain patterning media (**Fig. 2B, d7, triangles**). The embedding on d7 led to more pronounced bud extrusions 24 hours later (**Fig. 2B, d8, arrows**), before the EBs were transferred into spinning flasks bioreactors to promote the growth of the organoids (**Fig. 1D**). We quantified the area of EBs and hMOs, derived from five distinct iPSC lines, including two Parkinson’s disease lines, and compared their size from d2 to d8 within the disks, using an inhouse CellProfiler pipeline that created colored masks to visually verify the area to be quantified (**Fig. 2C**). Our analysis demonstrated that EBs from all lines grew in size from d2 to d8 within the disks, to attain a diameter of ~500-600 μm (**Fig. 2D**). The area of the hMOs generated at d8 from the disks were similar across all lines (**Fig. 2D**). After 10 days in spinning flasks, we counted hMOs from two different backgrounds, and counted 650 and 714 hMOs, respectively, in each of the bioreactors (**Fig. 3A**). The yield of material obtained from a 948-well EB-Disk was approximately ~65-70%. We noted the fusion of some hMOs that accounted for numbers being lower than our starting number of 948, as well as some hMOs being lodged in the bottom of the spinner impeller. Within each cell line, we also noted a homogenous size for the organoids when we collected and fixed our hMOs at 100 days (**Fig. 3B**).

To assess the cellular composition of the hMOs, we performed flow cytometry analysis with hMOs from our control line that was dissociated into single cell suspensions after 50 days in culture. The numbers of cells per hMO was on average ~106,674+/-SEM 7401 cells/hMO across two batches, and the viability was ~81.3% (**Fig. 4A**). Additionally, to assess the diversity of cell populations within the hMOs, we stained cryosections for S100β, MAP2 and TH, markers for astrocytes, neurons and a catecholaminergic neuron/rate-limiting enzyme in dopamine biosynthesis, respectively (**Fig. 4B**). Higher magnification revealed typical astrocytes, neurons and dopaminergic neurons morphologies (**Fig. 4C**). In 50-day old hMOs, in addition to immunocytochemistry, we utilized quantitative PCR (qPCR) to detect several transcripts known to be expressed in midbrain DNs, including nuclear receptor related 1 protein (Nurr1), LIM homeobox transcription factor 1-beta and alpha (LMX1A, LMX1B), TH, catechol-O-methyltransferase (COMT) and dopa decarboxylase (DDC). As expected, the expression of these markers was absent in the parental iPSCs (**Fig. 4D, red heatmap**). Conversely, OCT3/4 and NANOG, transcripts known to be expressed in iPSC were only detected in iPSCs and absent from the hMOs samples (**Fig. 4D, blue heatmap**). Additionally, after 100 days in final differentiation media, hMOs started to develop and accumulate spontaneous neuromelanin granules (**Fig. 4E**). This observation was previously confirmed to be a feature of functional dopaminergic neurons [7, 14, 63].

Next, to further validate the midbrain identity of our hMOs and the reproducibility of our method, we cleared and stained four 6-month-old hMOs from our control line for TH and MAP2. Consistent with published findings [7, 9, 14], we observed the presence of dopaminergic neurons stained for TH within the hMOs (**Fig. 5, thumbnails and Movie**). The volumetric quantification of MAP2 and TH staining percentages showed that similar levels of neurons and dopaminergic neurons were present across each of the four hMOs tested. The TH/MAP2 ratio showed a similar yields (~0.5) of dopaminergic neurons in four hMOs (**Fig. 5B**). Taken together, our findings indicate that large numbers of hMOs generated with this protocol present with good cell viability, a midbrain identity and similar cell types relative to each other and in line with earlier hMO protocols

**Fig. 5.**
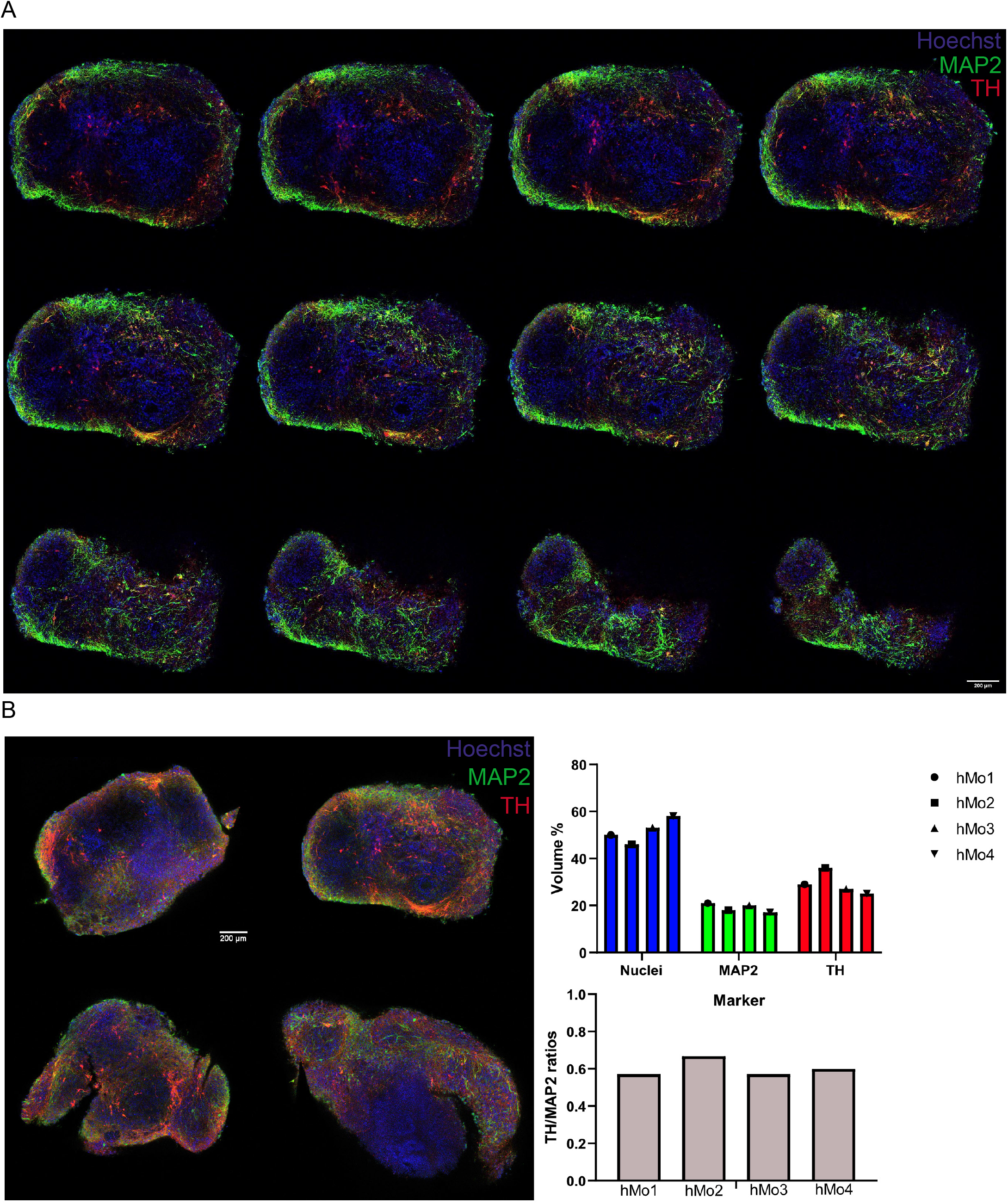
3D acquisition of hMOs images. (A) Thumbnails of 6-month old hMOs from Healthy line, cleared, stained, and imaged for MAP2 (green), TH (red), and nuclei (blue). Scale bar= 200 μm. See Movie. (B) Relative volumes of TH, MAP2 and nuclei quantifications and calculated TH/MAP2 ratios in four independent 6-month old hMOs from Healthy line.

## 4. Conclusion

In this manuscript, we describe a novel protocol applying a microfabricated EB-Disk device for the generate large batches of homogenous hMOs across multiple different iPSC lines. We describe in detail the critical points to successfully achieve hMO generation. For instance, the quality of iPSC colonies is the first critical aspect in generating high quality hMOs [7]. Our protocol presents the following advantages to users. Working with this method, they will be able to: 1) generate hundreds of hMOs for a given batch all from a single iPSC in one workflow; 2) avoid direct contact with EBs and minimize manual handling of hMO cultures; 3) lower reagent volumes required, by reducing the amount of media used for the generation and maintenance of hMOs, and also the quantity of Matrigel® reduced growth factor required for the embedding step, compared to previously described protocols [7, 14, 15].; and 4) reduce the overall cost of hMO production, by reducing the media changes to once a week, compared to three times a week as in our previous methods [7]. To note, this method does not require any robotic or automation system as described in other publications [30], and therefore can be used across the majority of laboratories, once they have access to an incubator, a magnetic stir plate and several glass bioreactors, equipment that can be easily purchased.

Upscaling has been a focus of the organoid field and different methods are being explored to achieve a higher throughput. Spinning bioreactors are used to grow large numbers of organoids and have the benefit of providing improved oxygen and nutrient distribution to the organoids in long term culture [15]. Regarding brain organoids, spinning bioreactors have been used by several groups [15, 36, 64], and while they require a large amount of media and space, maintenance is less time consuming and does not require manipulation to the same extent as multi-well plates. Qian and colleagues [17] developed miniaturized bioreactors, reducing considerably the amount of media needed. However, this required 3D printing and a complex assembly, and does not reduce the need for manual handling of the culture vessels. Other types of organoids can also be created using adapted protocols with the EB-Disk to model diseases that affect other areas of the brain, or even other organs, opening up a number of new possibilities for these microfabricated devices. Depending on the organoid type, stem cells can be cultured in a different vessel prior to transfer into spinner flask bioreactors or can be directly seeded into them. Groups with an interest in tissue transplantation and regenerative medicine successfully upscaled their organoid production for liver or pancreas, by directly seeding cells in the flasks which permits an increase in the size of their organoids and the number of organoids in culture at a time [65, 66].

In this manuscript, while confirming the midbrain identity, we showed specific cell populations were present within the hMOs (astrocytes, dopaminergic neurons and MAP2-positive neurons). A deeper characterization was performed previously [7] by single cell RNA sequencing of hMOs produced using the same timeline and patterning factors as described here. In our previous paper, we showed the presence of diverse cell populations at 50 days, including radial glia, astrocytes, and several different types of neurons, including dopaminergic neurons characterized by broader dopaminergic markers. Overall, this method manuscript is aimed at helping the scientific community to more efficiently generate hMOs for Parkinson’s disease studies.

## CRediT authorship contribution statement

**Nguyen-Vi Mohamed:** Conceptualization; Data curation; Formal analysis; Funding acquisition; Investigation; Methodology; Writing - original draft. **Paula Lépine:** Methodology; Writing - original draft. **María Lacalle-Aurioles**: Conceptualization; Data curation; Formal analysis; Methodology; Writing - original draft. **Julien Sirois:** Data curation; Formal analysis; Methodology. **Meghna Mathur:** Methodology. **Wolfgang Reintsch:** Conceptualization; Methodology. **Lenore K. Beitel:** Writing - review & editing. **Edward A. Fon:** Funding acquisition; Project administration; Resources; Supervision; Validation. **Thomas M. Durcan:** Funding acquisition; Project administration; Resources; Supervision; Validation; Writing - review & editing.

## Acknowledgements

TMD and EAF received funding to support this project from the Canada First Research Excellence Fund, awarded through the Healthy Brains, Healthy Lives initiative at McGill University, the CQDM Quantum Leaps program and the Sebastian and Ghislaine Van Berkom Foundation. EAF is supported by a Foundation grant from CIHR (FDN-154301), a Canada Research Chair (Tier 1) in Parkinson Disease and the Canadian Consortium on Neurodegeneration in Aging (CCNA). TMD is supported by a project grant from CIHR (PJT - 169095). TMD also received funding support for this project through the Ellen Foundation. MNV is supported by a FRSQ and Parkinson Canada postdoctoral fellowship.

We acknowledge Dr. Tilo Kunath and Dr. Karamjit Singh Dolt (The University of Edinburgh) for providing the PD iPSC lines (AST23, AST23-2KO, AST23-4KO) and the C-BIG Biorepository Histology Core Facility (C-BIG) for the PD 2 and control iPSC lines. We also wish to thank Dr. Chen Carol X.-Q. and Dr. Narges Abdian for the quality control of iPSC lines used, Leonardo La Calle for editing the 6-month-old hMO movie, and acknowledge the MNI Microscopy Core Facility for confocal microscope management and maintenance.

---

**Fig. S1.**
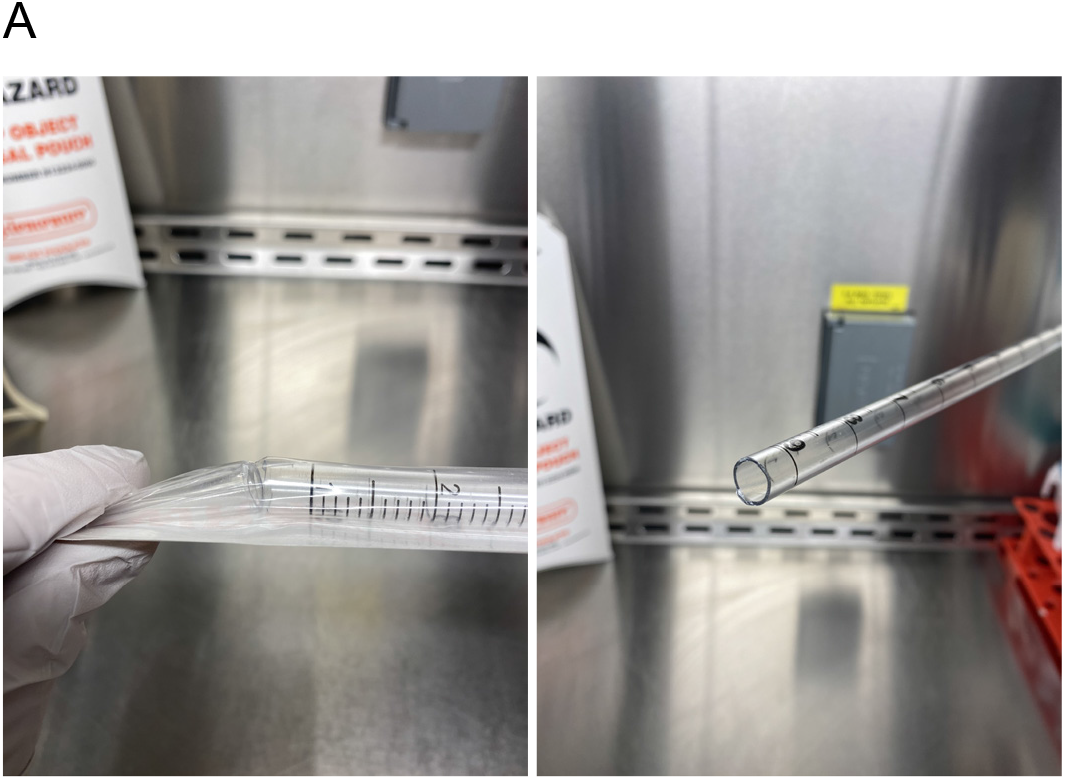
Supplemental method information (A) Representative picture of breaking the tip off of a 10 mL pipet under the biosafety cabinet, in sterile conditions.

## Abbreviations

(iPSCs): induced pluripotent stem cells
(PD): Parkinson’s disease
(DA): dopaminergic neurons
(hMOs): human midbrain organoids
(EB disk): embryoid-bodies disk device

